# Engineered spermidine-secreting *Saccharomyces boulardii* ameliorate colitis and colon cancer in mice

**DOI:** 10.1101/2025.01.19.633601

**Authors:** Mahdi Mohaqiq, Roger Palou, Ruizhen Li, Guijun Zhang, Het Vaishnav, Florance Parweez, Terence Moyana, Damian Carragher, D.W. Cameron, Timothy Ramsay, Almer van der Sloot, Sanjay Murthy, Brian Coombes, Mike Tyers, X. Johné Liu

## Abstract

Experimental studies suggest that the probiotic yeast *Saccharomyces boulardii* can mitigate the symptoms of inflammatory bowel disease. However, these results are equivocal and *S boulardii* probiotic therapy has not gained widespread acceptance in clinical practice. To assess whether the therapeutic properties of *S boulardii* might be improved upon, we engineered *S boulardii* to overproduce and secrete spermidine, a pro-regenerative natural metabolite. We employed CRISPR gene deletion and transposon-mediated gene integration to manipulate expression of key enzymes in the polyamine synthetic and transport pathways. We tested the engineered yeast by oral gavage of mice treated with azoxymethane and dextran sulfate sodium to induce chronic colitis and colon cancer. We demonstrate that oral delivery of spermidine-secreting *S boulardii* in mice populates the gastrointestinal tract with viable spermidine-secreting *S boulardii* cells and raises free spermidine levels in the gastrointestinal tract. Strikingly, spermidine-secreting *S boulardii* strains were significantly more effective than wild-type *S boulardii* in reducing dextran sulfate sodium-induced colitis as well as colitis-associated colon cancer in mice. These results suggest that in situ spermidine secretion by engineered synthetic biotic yeast strains may be an effective and low-cost therapy to mitigate inflammatory bowel disease and colon cancer.

## Introduction

Inflammatory bowel disease (IBD), including Crohn’s disease and ulcerative colitis (UC), is a highly morbid disease affecting the intestines and extra-intestinal organs. IBD is associated with substantially increased risk of hospitalizations, intestinal surgeries ^1, 2^ and cancer ^3^. Canada has among the highest prevalence of IBD in high income countries, with an estimated 825 per 100,000 people (>320,000 in the total population) living with the disease ^3^. The incidence of IBD is rapidly rising in developing low- and middle-income, while the prevalence also continues to rise further in high income countries due to an aging demographic ^4^.

The pathogenesis of IBD involves an aberrant immune reaction to commensal gut flora in genetically susceptible individuals, precipitated by a leaky gut barrier that permits passage of mucosal bacteria into the deeper layers of the bowel wall ^5, 6^. The resulting inflammatory reaction damages the gut lining through widespread activation of effector T cells and massive release of cytokines such as tumor necrosis factor α (TNFα) and inflammatory interleukins (IL)^7^. Numerous studies have also demonstrated microbiome dysbiosis in the gastrointestinal tract (GI tract) of individuals with IBD, particularly in the presence of gut inflammation, with reduced diversity of bacterial species as a hallmark feature ^8^. Notwithstanding, there are limited data demonstrating effectiveness of microbiome-modulating therapies, such as restrictive diets, prebiotics, probiotics, antibiotics, or fecal microbiota transplant ^9, 10^.

Recent IBD treatments directly target the immune system. Over the past two decades, there has been gradual introduction and adoption of multiple classes of biologic and small molecule targeted therapies to treat IBD ^9, 11^. While these treatments have generally shown greater effectiveness than conventional therapies, less than 50% of IBD patients achieve lasting disease control even with maximal therapeutic doses, underscoring the challenges in treating this chronic disabling condition ^12^. As such, IBD patients continue to face high rates of morbidity, disability, and lost productivity ^13–16^. At the same time, many of the systemic immunosuppressive therapies used to treat IBD are associated with increased risks of adverse reactions, most notably serious infections ^17^ and cancers ^18^. Furthermore, these newer treatment options come at substantial costs, placing considerable strain on health care resources. Targeted therapies now account for the greatest proportion of health care spending in IBD patients ^16, 19^. As a general class, biologics to treat immune mediated diseases account for close to 30% of pharmaceutical health care spending in Canada ^20^.

Chronic bowel inflammation is a major risk factor for colorectal cancer (CRC). IBD patients carry a 2-fold increased lifetime risk of developing CRC relative to age-sex-matched individuals without IBD and a 5-10% absolute lifetime risk ^21^. This risk is exacerbated by higher cumulative lifetime colorectal inflammatory burden, primary sclerosing cholangitis, a personal history of colorectal dysplasia, and/or a first-degree relative diagnosed with CRC before the age of 50 ^22–25^. Furthermore, IBD-CRC has poorer prognosis than sporadic CRC due to its distinct characteristics such as younger age at diagnosis, diagnosis as an emergency, an increased likelihood of right-sided colon involvement, and histological features like multifocal tumors, poor/undifferentiated histology, or mucinous carcinomas ^26^. Given the shortcomings of available therapies to treat IBD, including limited efficacy, potential safety concerns and high costs, targeted delivery of an efficacious, cost-effective therapy to the gut to a achieve sustained disease remission would be transformative for IBD patients.

Several pilot studies have suggested that the probiotic yeast species *S boulardii*, either alone ^27^ or in combination with mesalazine ^28^, can be effective in treating UC patients who are either intolerant or refractory to mesalazine therapy. However, no follow-up randomized control trials have been published using *S boulardii* in UC and one trial in Crohn’s disease showed no benefits^29^. Therefore, *S boulardii* is not used as a standard therapy to treat IBD in clinical practice ^30, 31^.

Nonetheless, the *S boulardii* holds promise as living therapeutics for in situ production and delivery of biologics in IBD because of its inherent mild anti-inflammatory effects, ability to transiently colonize the gut mucosa, and established safety profile. *S boulardii* has an additional advantage over more established bacterial synthetic biotics ^32, 33^ both because of its compatibility with antibiotics that are often used in the management of IBD ^34^ and minimal risk of lateral gene transfer to other species in the microbiome ^35^.

To engineer an effective IBD therapeutic, we focused on the polyamine spermidine. First, intestinal mucosa-derived spermidine is vital for mucosa maturation in juvenile rats and in mucosal healing from cytosine-induced intestinal injury in adult rats ^36^. Targeting mucosal healing would address the greatest unmet need in IBD therapy ^37^. Second, dietary supplementation with high levels of exogenous spermidine reduces experimentally induced colitis in mice ^38, 39^. Here we report the generation of engineered *S boulardii* strains that overproduce and secrete spermidine to deliver spermidine directly to the gut mucosa. In vivo studies in mice with spermidine secreting *S boulardii* strains demonstrated striking efficacy in reducing dextran sulfate sodium (DSS)-induced chronic colitis and colitis associated colon cancer in mice ^40^.

## Results

### Engineering S. boulardii to over-produce and export spermidine

*S. boulardii* shares greater than 99% of its genome sequence^41^ with the bakers’ yeast *S. cerevisiae* which has well-developed molecular genetic tools as a result of decades of use as a major model organism in biomedical research. Despite its close relationship to *S cerevisiae*, however, *S boulardii* has not been widely used in biomedical research because of its relatively poor transformation efficiency, the absence of *S boulardii* strains bearing standard nutrient auxotrophies, and the lack of validated vector systems for transformation and gene expression control. *S cerevisiae* has previously been modified to overproduce and export polyamines for the purposes of high-level in vitro production of polyamines in biotechnological applications ^42^. For example, an *S cerevisiae* strain called OS123/pTPO1 has been engineered to overexpresses three polyamine biosynthesis genes (*SPE1*, *SPE2*, *SPE3*) and the polyamine transporter *TPO1*, in conjunction with a deletion in the *OAZ1* gene that encodes ODC antizyme (see Fig. 1A for summary of polyamine biosynthesis in yeast). However, Tpo1, and other members of this family (Tpo2-4) ^43^, are proton-polyamine antiporters that likely function to mediate polyamine uptake into the acidic yeast vacuole ^44^. Export of polyamines through plasma membrane localized Tpo1 thus requires an acidic extracellular pH ^42, 45^. Unlike the highly acidic stomach, the lower GI tract has a slightly alkaline pH ^46^ such that Tpo1 will function as a spermidine importer in this environment. In contrast, the polyamine transporter Tpo5 resides in post-Golgi secretory vesicles ^47^ and mediate polyamines export through exocytosis in a manner that is dependent on intracellular polyamine concentrations. A recent study has reported that *TPO5* overexpression in *S. cerevisiae* results in ∼10% increase of spermidine export in saturated yeast culture ^48^.

**Figure 1.**
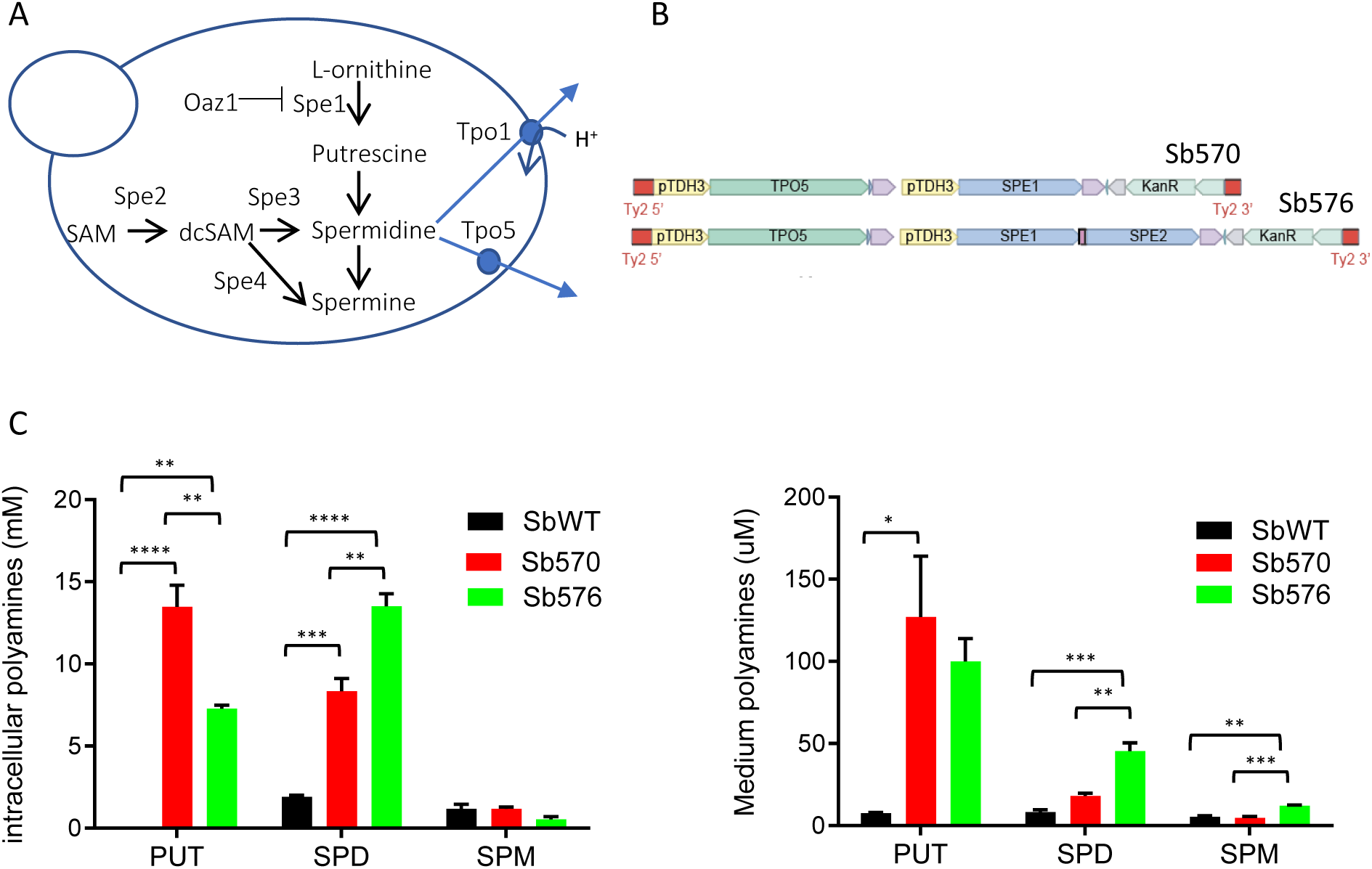
Engineering *S. boulardii* to overproduce and secrete spermidine. A. Polyamine synthetic pathway in *S cerevisiae*. See text for details. Tpo1 (pH dependent) and Tpo5 (pH-independent) are two possible polyamine exporters in *S cerevisiae* and *S boulardii*. B. Integration cassette for overexpression of *TPO5*, *SPE1* and *SPE2*. The Ty2 element is an LTR-containing retrotransposon present in several copies in the genomes of *S. cerevisiae* and *S. boulardii*. The TPO5, SPE1 and/or SPE2 genes were expressed from the *TDH3* promoter. The kanMX marker allows growth selection on G418 medium. 5’ and 3’ Ty2 recombination arms flank the entire cassette. C. Intracellular (left) and culture medium (right) polyamine concentrations in saturated yeast cultures grown in synthetic complete (SC) medium. Means ± SEM for three independent experiments are shown. PUT: putrescine; SPD: spermidine; SPM: spermine. Significance was calculated by One-way ANOVA and Tukey’s multiple comparisons test.

Based on the above studies, we sought to engineer an *S boulardii* strain that overexpresses both polyamine biosynthesis genes and the *TPO5* transporter gene. We optimized *S boulardii* transformation, genetic selection and CRISPR-based gene editing for strain construction (see Materials and Methods). To ensure genetic stability and homogenous gene expression in all cells, we employed transposon-mediated gene integration ^49, 50^ to overexpress polyamine biosynthesis genes and *TPO5* in *S boulardii*. A multi-gene encoding plasmid was integrated in the Ty2 locus ^41^ of the *S boulardii* parental strain MYA-796 to generate two *S boulardii* strains that overproduced and exported spermidine, Sb570 and Sb576 (Fig. 1B; genotypes of all *S boulardii* strains are listed in Supplementary Table 1). Sb570 harbored an *OAZ1* deletion by CRISPR-based gene editing and integrated exogenous copies of *SPE1* and *TPO5*, while Sb576 additionally harbored an integrated copy of *SPE2* at the Ty2 locus. To facilitate detection of the engineered *S boulardii* strains in the mouse GI tract, the integrative vectors also included a kanMX marker cassette. Both Sb570 and Sb576 exhibited growth rates similar to that of the wild type (WT) *S boulardii* strain (SbWT, data not shown). We measured intracellular (i.e., in the cell pellet) and secreted (i.e., in the culture medium) polyamine concentrations of saturated cultures of the two engineered yeast strains as compared to SbWT strain. These results confirmed that both strains produced and secreted significantly more spermidine than the parental strain (Fig. 1C).

### Spermidine secreting S boulardii elevated luminal spermidine levels in the GI tract

We focused on strain Sb576, which produced and secreted more spermidine than Sb570 (Fig. 1C), in our animal studies. We first determined if Sb576 retains viability in the GI tract. Female mice were treated with the antibiotic streptomycin to reduce the endogenous bacterial microbiome and facilitate yeast retention in the GI tract ^51, 52^. Following streptomycin treatment, mice received a daily oral gavage of vehicle control, SbWT or Sb576 for five days at a dose of 1 × 10^8^ yeast cells per mouse per day (Fig. 2A). Fresh fecal samples plated on medium containing ampicillin revealed similar levels of viable SbWT and Sb576 (Fig. 2B) demonstrating that spermidine production did not impact Sb576 competitive fitness in the mouse GI tract. We found no yeast colonies in serially diluted fecal samples from mice not gavaged with yeast, indicating that the endogenous yeast microbiome is relatively low (< 2×10^3^ CFU/g feces, compared to >2×10^6^ CFU/g feces in mice gavaged with yeast). As expected, while no yeast colonies were found in fecal samples from SbWT-treated mice on G418 plates (not shown), similar amounts of G418 resistant and ampicillin resistant colonies were found in fecal samples from Sb576-treated mice (Fig. 2B) since Sb576 contains an integrated KanMX marker (Fig. 1B). To assess whether the engineered Sb could elevate spermidine levels in the GI tract, we next compared free spermidine concentrations in fresh fecal samples of mice following the fifth oral gavage (on day 0 prior to AOM injection) (Fig. 2A). Administration of Sb576, but not SbWT, significantly increased fecal spermidine levels compared to vehicle-treated mice (Fig. 2C).

**Figure 2.**
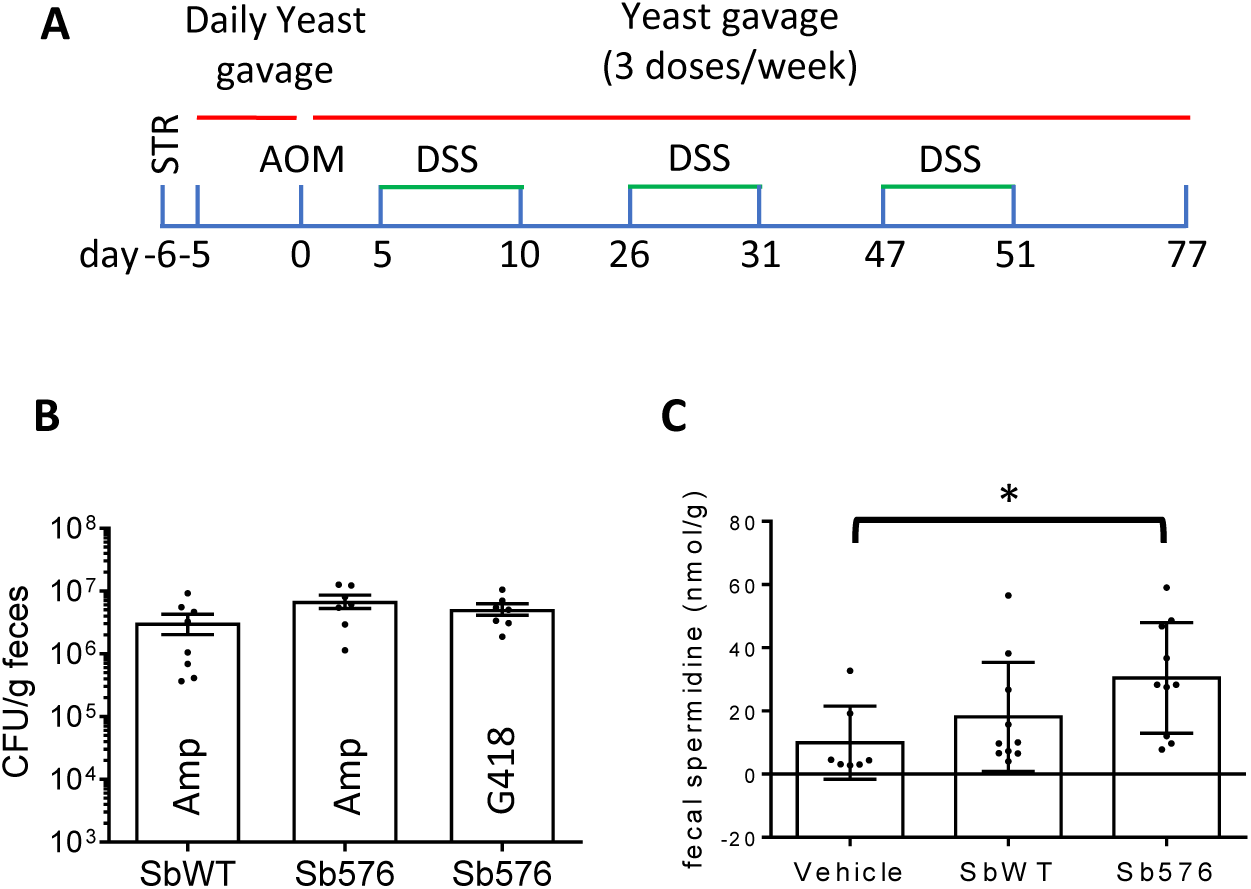
Sb576 increases colonic spermidine. A. Schematic of AOM-DSS protocol (See text for details). B. Yeast CFU after five days of oral gavage with SbWT or Sb576 (both at 1×10^8^ cells per mouse per day). Fecal samples from Sb576-treated mice were plated on both ampicillin- and G418-containing plates. Shown are means ± SEM (CFU per g fresh fecal sample). Each dot represents one mouse. C. Free spermidine levels in fresh fecal samples following five days of oral gavage of vehicle, SbWT or Sb576. Shown are means ± SEM. One-way ANOVA and Tukey’s multiple comparisons test.

### Spermidine secreting S boulardii reduces colitis and colitis associated colon cancer in mice

Following the initial five daily oral gavages of vehicle or yeast, mice then received a single injection (i.p.) of azoxymethane (AOM, 12.5mg/kg) followed by three sequential cycles of DSS treatment. Vehicle or yeast oral gavage was continued three times a week after the first five days (Fig. 2A). We monitored daily body weight changes (except for control mice which were monitored intermittently), as well as stool consistency and blood in stools, the three components of colitis disease activity index (DAI, see Materials and Methods for details) for 10 days in each DSS treatment cycle. As shown in Figure 3A, all three groups of mice treated with AOM and DSS lost considerable weight compared to untreated control mice. However, Sb576 treatment, but not SbWT treatment, significantly reduced body weight loss compared to vehicle treatment. Furthermore, Sb576 treatment significantly reduced colitis disease burden (i.e. DAI) compared to SbWT or vehicle treatment, although SbWT treatment also significantly reduced DAI compared to vehicle control (Fig. 3B). As another hallmark of DSS-induced colitis, AOM/DSS-treated mice had significantly shorten colon (Fig. 4A, control vs. vehicle groups). SbWT treatment moderately, but significantly, reduced colon shortening compared to vehicle treatment. However, colon lengths of mice treated with Sb576 were significantly greater than those of vehicle- or SbWT-treated mice. In fact, Sb576 treatment restored colon length to normal range of control mice (Fig. 4A). DSS-treatment also causes enlargement of the spleen in mice ^53^, although splenomegaly is not a common symptom of ulcerative colitis in humans ^54^. We confirmed spleen enlargement in DSS-treated mice (Fig. 4B, control vs. vehicle). However, we observed no significant changes in spleen weight comparing Sb576-treated mice with vehicle- or SbWT-treated mice (Fig. 4B). Together, these results (Figures 3A-B and 4A) demonstrate that Sb576 is significantly more effective than SbWT in ameliorating DSS-induced colitis in mice.

**Figure 3.**
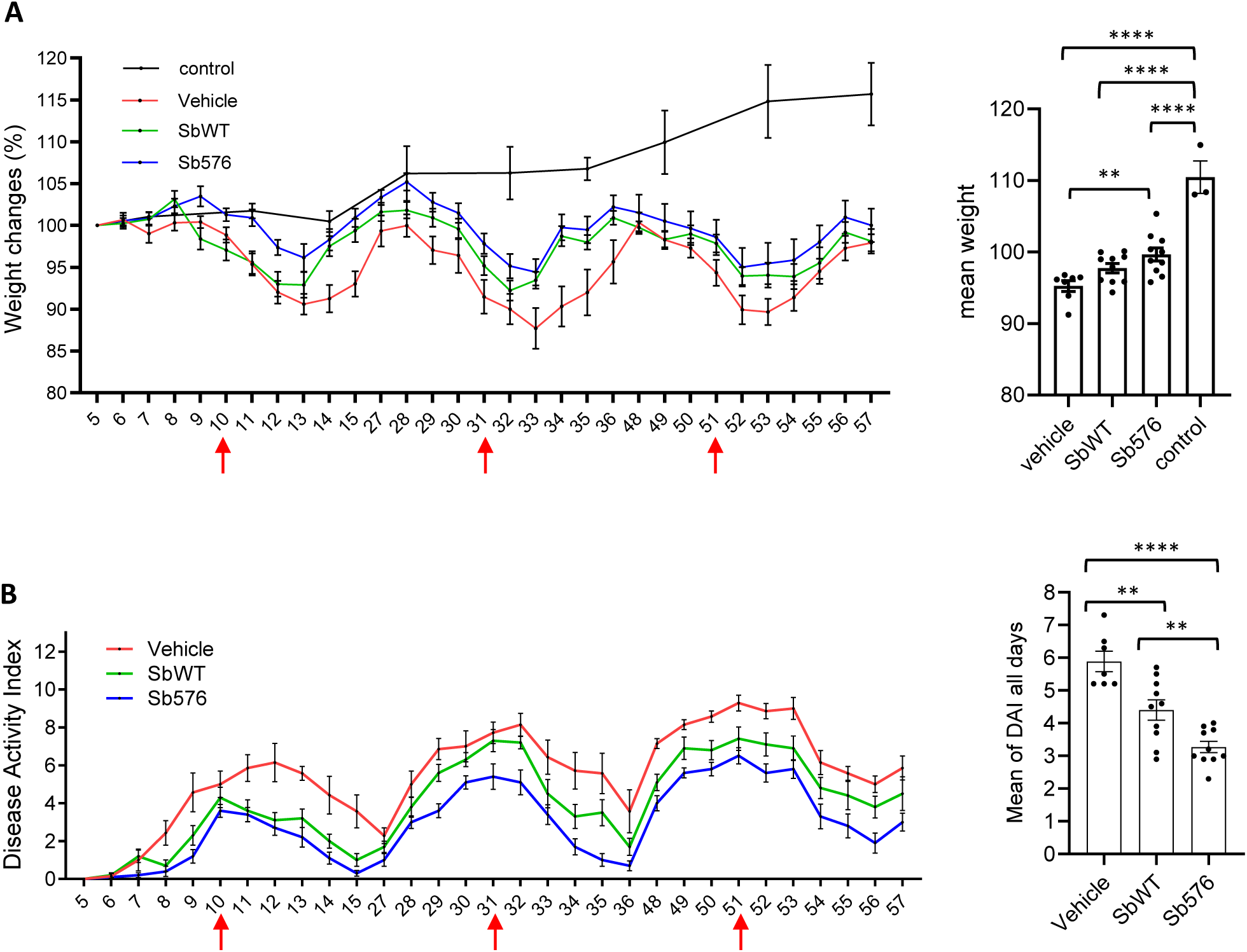
Sb576 reduces DSS-induced colitis symptoms. A. Daily body weight (means ± SEM) expressed as % of the weight on day 5 (prior to the first DSS cycle) of mice receiving vehicle (n=7), SbWT (n=10) or Sb576 (n=10), 10 consecutive days for each DSS cycle starting one day after the beginning of DSS treatment. Body weight of control mice (n=3) from the same cohort is shown on intermittent days. Shown on the right are mean body weights (means ± SEM) of individual mice over the 30-day course. One-way ANOVA and Tukey’s multiple comparisons test. Arrows indicate cessation of DSS water. B. Daily DAI (means ± SEM) of mice receiving vehicle (n=7), SbWT (n=10) or Sb576 (n=10) following each of the three DSS cycles, as above. Shown on the right are mean DAIs (means ± SEM) of individual mice over the 30-day course. One-way ANOVA and Tukey’s multiple comparisons test. Arrows indicate cessation of DSS water.

**Figure 4.**
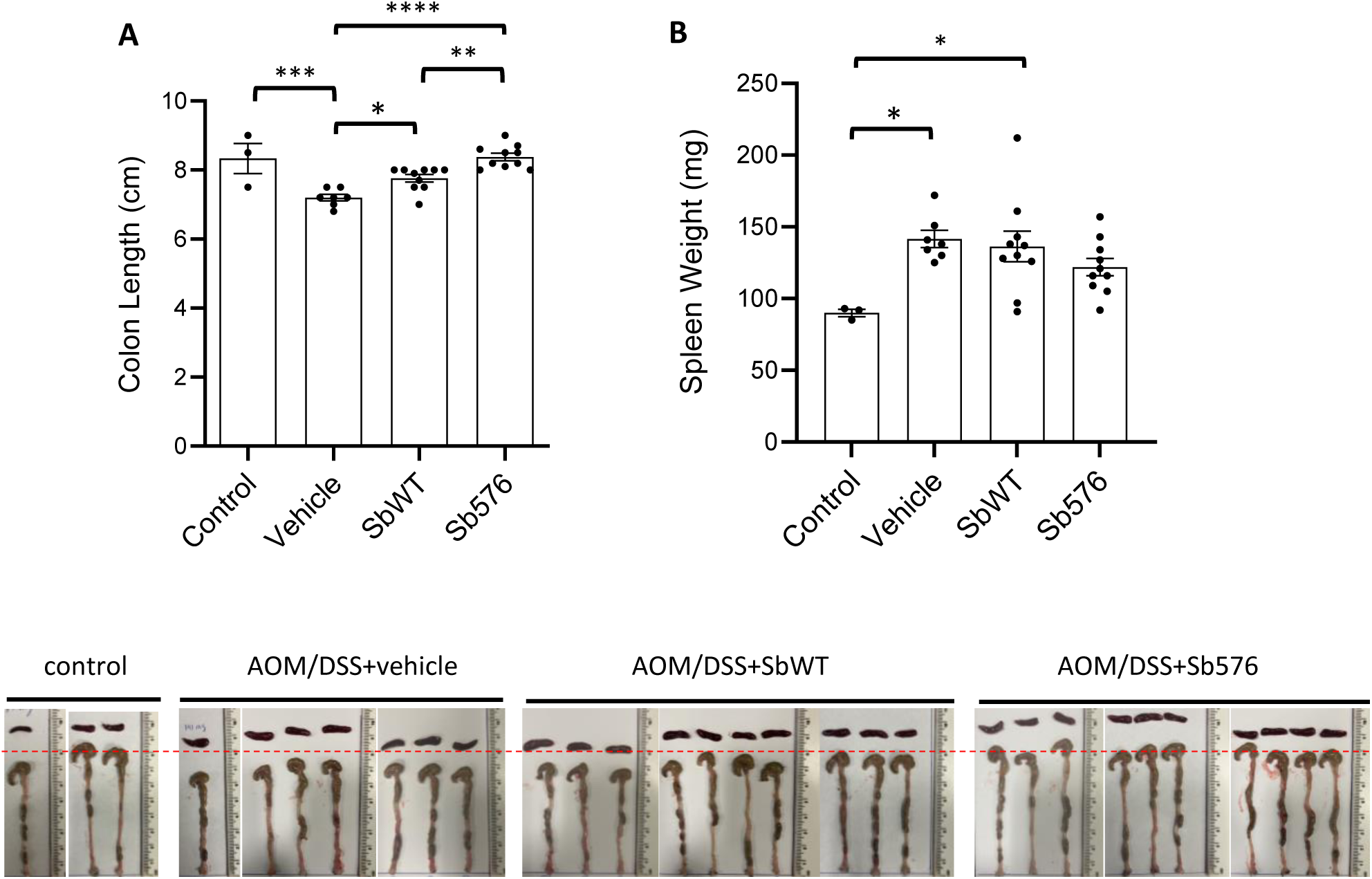
Sb576 restores colon length in AOM/DSS-treated mice. A. Colon length of the four groups of mice. Shown are means ± SEM. One-way ANOVA and Tukey’s multiple comparisons test. B. Spleen weight of the four groups of mice. Shown are means ± SEM. One-way ANOVA and Tukey’s multiple comparisons test. Colon and spleen images of the entire cohort of mice are shown below. A dashed line is drawn across the 8 cm marks of the measuring tape.

To investigate the effect of Sb576 on the AOM/DSS-induced colon cancer phenotype, we assessed colon tumor numbers and tumor burden (sum of tumor area, in mm^2^) ^39^. Whereas no tumors were found in control mice (Fig. 5A, control image), AOM/DSS mice treated with vehicle alone had numerous tumors, mainly found in the middle and distal colon (Fig. 5A, vehicle image; quantitative data shown in Fig. 5B and 5C), consistent with previously published study ^40^. We found no visible tumors in any other major organs in AOM/DSS-treated mice (not shown). AOM/DSS-treated mice gavaged with SbWT or Sb576 had significantly fewer numbers (Fig. 5B) and lower tumor burden (Fig. 5C) than AOM/DSS-treated mice gavaged with vehicle. The difference between Sb576 and SbWT treatment was not significant.

**Figure 5.**
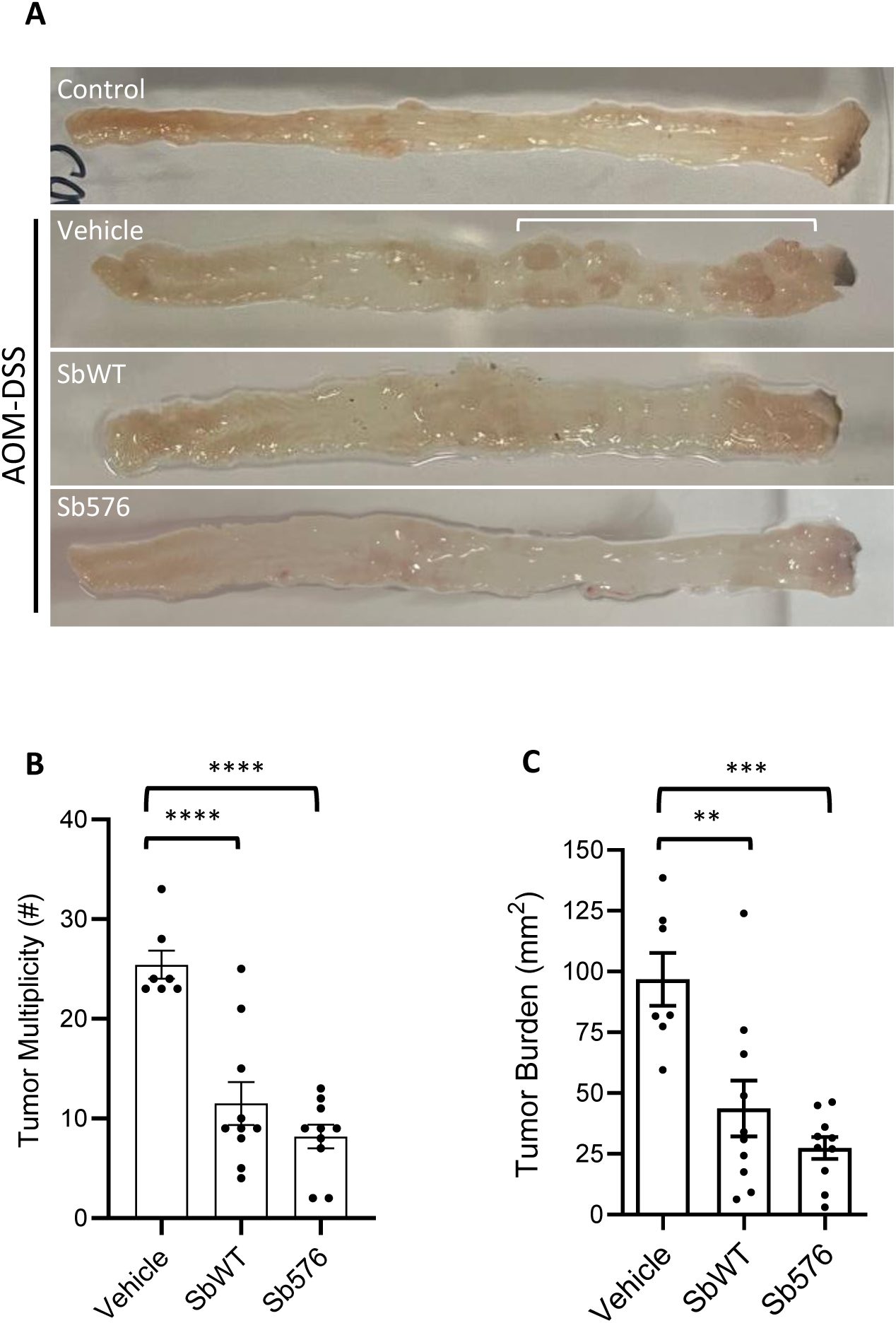
Sb576 reduces macroscopic colon tumors. A. Representative colon images (taken by cellphone camera during tumor counting/measuring) of untreated control, AOM-DSS-treated mice receiving vehicle, SbWT or Sb576. Distal ends on the right. White line marks tumors in the colon of AOM-DSS mice treated with vehicle. Please note that tumor counting and measuring were done under a dissecting microscope. B. Tumor numbers. C. Tumor burden (sum of tumor area). Shown are means ± SEM. One-way ANOVA and Tukey’s multiple comparisons test. Below: Colon images of representative mice of the four groups taken by a cellphone camera. Insets show the respective images taken (by a cellphone camera) through the eye piece of a dissecting microscope while measuring tumors.

Immediately following macroscopic tumor scoring (above), the colons were fixed as a Swiss roll for further histopathology analyses to assess microscopic neoplastic lesions. A single longitudinal paraffin section through the center of the colon (see examples in Fig. 6, control and AOM-DSS) was examined microscopically to classify all neoplastic lesions ^40, 55–57^. Whereas colonic sections from control mice exhibited the normal mucosal morphology without any neoplastic lesions, sections from AOM-DSS-treated mice exhibited various neoplastic changes, classified as low-grade adenomas, high-grade adenomas, or adenocarcinomas (see Materials and Methods for details). Our data indicated that while there was no significant difference in the numbers of low-grade adenomas among the vehicle only, SbWT and Sb576 groups, AOM/DSS mice gavaged with either SbWT or Sb576 had significantly fewer high-grade adenomas and adenocarcinomas compared to vehicle only (Fig. 6). Notably, animals treated with Sb576 had significantly fewer total neoplastic lesions than either of the other two groups (Fig. 6). Taken together, these data demonstrated that both SbWT and Sb576 were able to significantly reduce colon cancer progression in the AOM/DSS mouse model, and that Sb576 was significantly more effective than SbWT in doing so.

**Figure 6.**
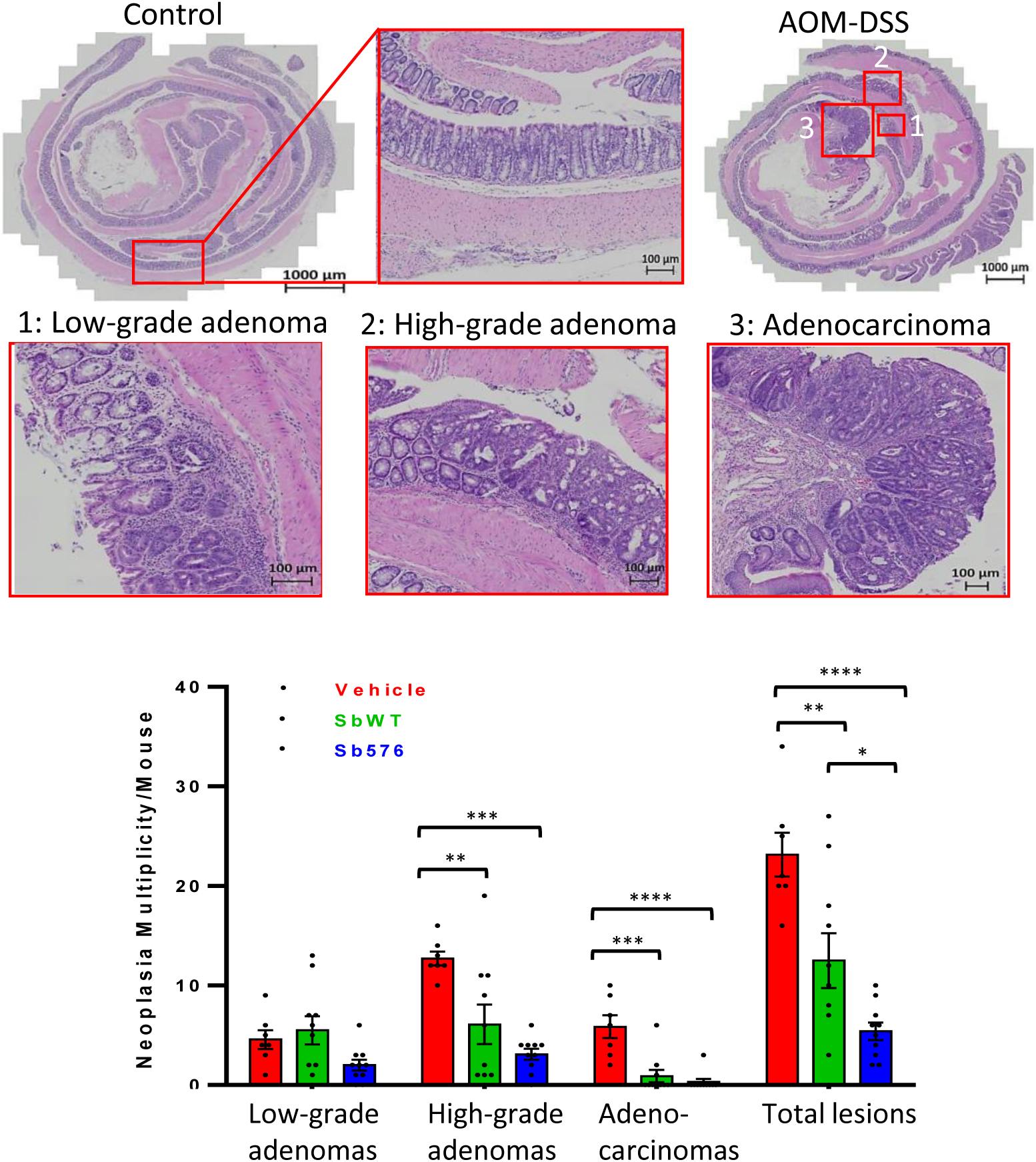
Sb576 reduces microscopic neoplastic lesions. Hematoxylin and Eosin staining images of colonic neoplasia. Control: a section through the middle of the colon Swiss roll (distal end at center) of a control mouse showing normal colonic crypts. AOM-DSS: A colon Swiss roll section of a mouse in the AOM/DSS-vehicle group depicting low-grade and high-grade adenomas, and adenocarcinoma. The graph summarizes the numbers of low-grade or high-grade adenomas, adenocarcinomas and total neoplastic lesions in AOM-DSS treated mice receiving vehicle, SbWT or Sb576 treatment. Shown are means ± SEM (n=7 for vehicle; n=10 for SbWT or Sb576). One-way ANOVA and Tukey’s multiple comparisons test.

### Spermidine secreting S boulardii does not activate cytotoxic T cells

Since gut inflammation (i.e., colitis) is a key factor for colon cancer development in the AOM/DSS mouse model ^58^, the ability of Sb576 to reduce tumor burden and neoplastic lesions is likely attributable to its ability to reduce colitis. However, it is conceivable that in situ spermidine production by Sb576 may also reduce tumor development by other mechanisms, for example by enhancing host antitumor immunity ^59^. To examine this latter possibility, we assessed cytotoxic T cells associated with neoplastic lesions using anti-CD8 and anti-Granzyme B immunohistochemistry of adjacent colon sections (see Fig. 7A, left and right, respectively). We noted a significant increase of reactive lymphoid follicles (RLF) in mice treated with AOM/DSS and gavaged with vehicle only compared to control untreated mice (Fig. 7B, left). AOM-DSS mice treated with SbWT or Sb576 had significantly reduced number of RLFs compared to vehicle only group, although there was no significant difference between SbWT and Sb576 (Fig. 7B, left). In contrast, there was no significant difference in the proportion of CD8 positive T cells in the RLFs between any groups (Fig. 7B, right). The CD8 positive T cells in RLFs were mostly negative for Granzyme B (Fig. 7A, second row), suggesting that these were naïve T cells. Despite the significant difference in total neoplastic lesion counts between vehicle only, SbWT and Sb576 treated animals (Fig. 6), we found no significant difference in the proportions of CD8 positive T cells (Fig. 7C, left), or Granzyme B positive cells (Fig. 7C, right), associated with the neoplastic lesions in the three treatment groups. These results suggested that the increased numbers of RLFs observed in the vehicle only treatment group are likely the result of increased inflammation ^60^. Our results also suggest that the reduced tumor burden in SbWT and or Sb576 treated groups may be independent of cytotoxic CD8^+^ T-cells.

**Figure 7.**
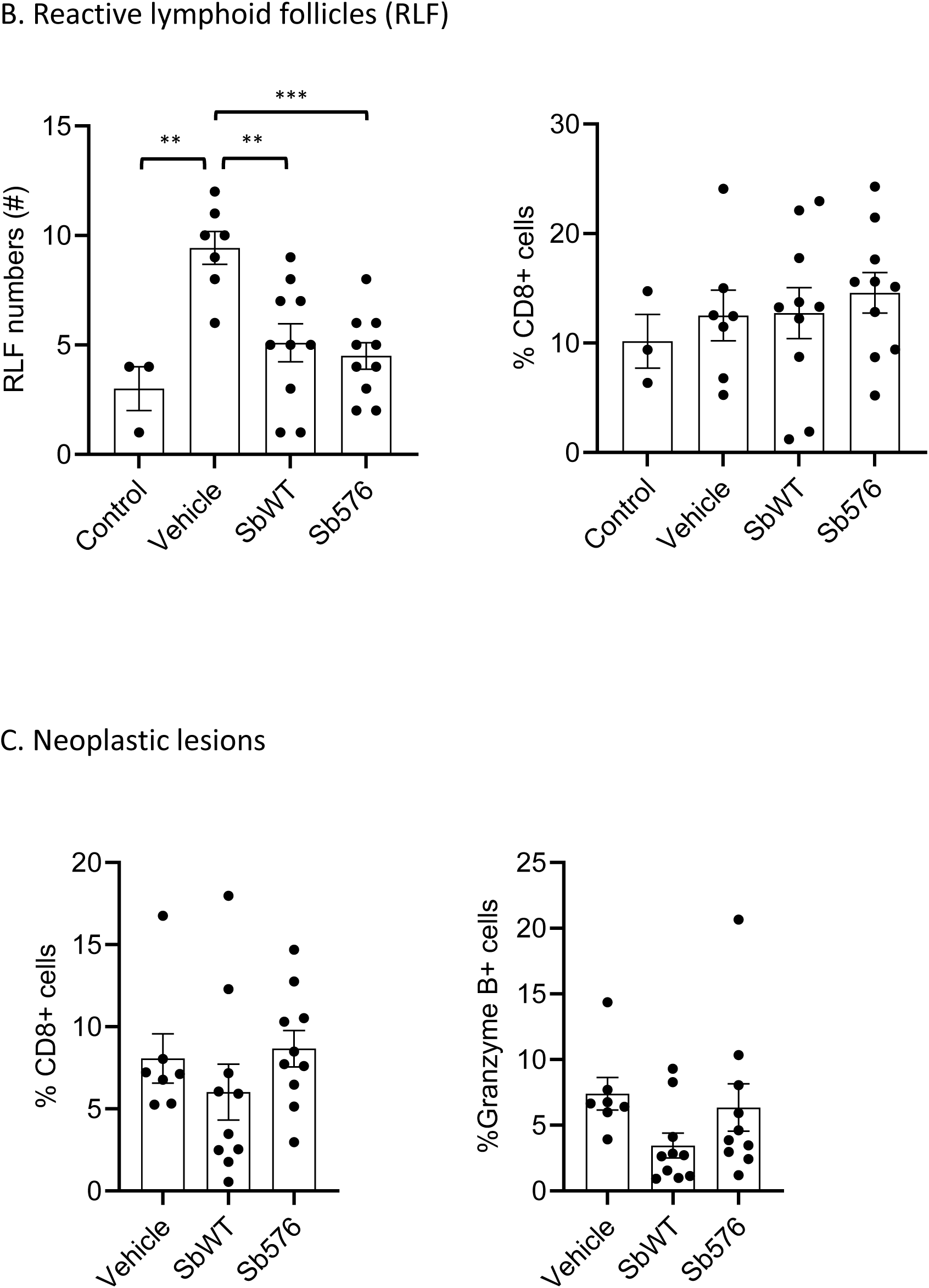
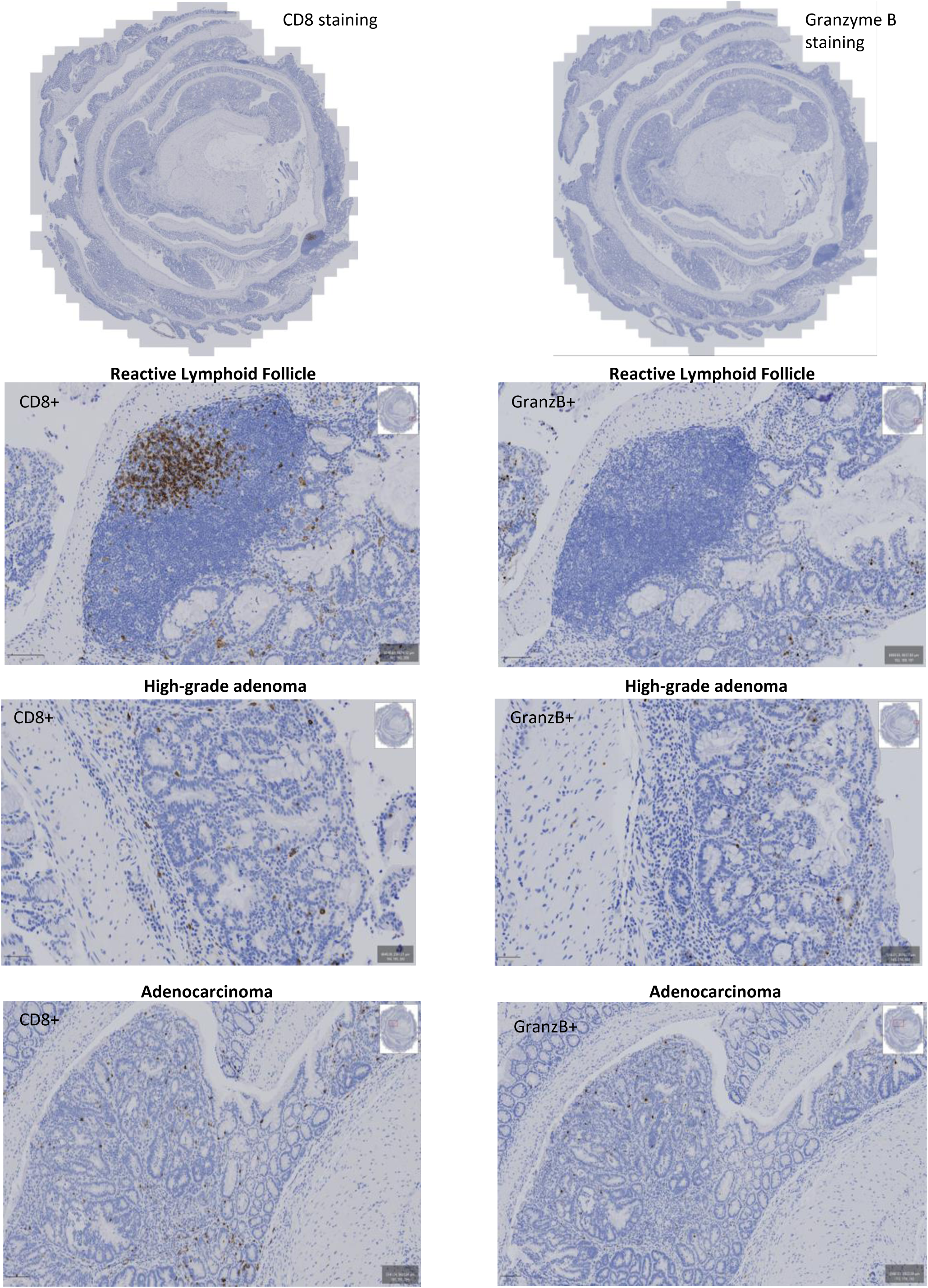
Sb576 does not activate CD8 lymphocytes. A. Anti-CD8 (left) and anti-Granzyme B (right) immunohistochemistry of adjacent sections of an AOM-DSS mouse colon. A reactive lymphoid follicle, a high-grade adenoma and an adenocarcinoma are highlighted. B. Reactive lymphoid follicle (RLF) counts (left) and proportions of CD8^+^ T cells within the RLF (right). Shown are means ± SEM. One-way ANOVA and Tukey’s multiple comparisons test. C. Proportions of CD8^+^ T cells (left) and proportions of Granzyme B positive cells (right) in neoplastic lesions of AOM-DSS mice treated with vehicle, SbWT or Sb576. Shown are means ± SEM.

## Discussion

We have engineered *S. boulardii*, a U.S. Food and Drug Administration (FDA)-designated Generally Recognized as Safe (GRAS) organism, to overproduce and secrete spermidine. We have demonstrated that the spermidine secreting *S boulardii* strain, Sb576, retains viability in the mouse gut and increases free spermidine levels in the gut lumen. Importantly, Sb576 significantly reduced colitis symptoms and colitis-associated colon carcinogenesis in mice.

Sb576 exhibited significantly greater efficacy compared to SbWT in reducing colitis symptoms including weight loss and DAI. Most significantly, Sb576-treatment completely restored colon length to normal range of untreated mice. Both SbWT and Sb576 significantly reduced colitis-associated colon carcinogenesis compared to vehicle-treated mice. Although macroscopic tumor burden between mice treated with Sb576 and SbWT was not significantly different, mice treated with Sb576 exhibited significantly fewer total neoplastic lesions than mice treated with SbWT. Therefore, Sb576 was also significantly more effective than SbWT in reducing AOM/DSS-induced colon carcinogenesis.

Consistent with our results, it has been reported that wild type but not polyamine deficient *E. coli* can increase gut epithelial cell polyamine concentrations and reduce DSS colitis in a gnotobiotic mouse model ^61^. We note, however, that our data were obtained in the context of complex natural microbiome and thus directly extensible to human disease contexts. To our knowledge, the results presented here are the first to demonstrate efficacy of an engineered probiotic organism that secretes spermidine to ameliorate colonic inflammation.

In addition to ameliorating colitis in mice ^38, 39^, spermidine supplementation has shown effectiveness in promoting longevity and in mitigating various other disease phenotypes ^62^. A recent study suggests that fasting-mediated autophagy and longevity requires endogenous polyamine, especially spermidine, synthesis ^63^. However, most animal studies have employed high doses of exogenous spermidine, which may carry long-term health risks. For instance, some cancers and stroke have been associated with increased polyamine metabolism ^64, 65^ and tumor cell-derived spermidine may suppress intratumoral CD8^+^ T cell activation ^66^. The approach of delivering spermidine in situ using engineered *S. boulardii* strains may circumvent the drawbacks of systemic oral supplementation by producing spermidine locally, where appropriate interaction with gut epithelium, the associated immune system ^67, 68^ and endocrine system ^68^ can occur. This may be particularly relevant in treating inflammatory bowel disease given the critical role of mucosal polyamines, especially putrescine and spermidine, in intestinal mucosal recovery from chemotherapy-induced injury in adult rats reported decades ago ^36^. In providing in situ synthesis and secretion of spermidine, Sb576 may recapitulate the regenerative prowess of natural polyamines and promote mucosal healing in mice subjected to multiple cycles of DSS treatment. Indeed, despite significant gut inflammation, as evident by body weight loss, DAI score and colon carcinogenesis, AOM/DSS mice treated with Sb576 had colon length indistinguishable from those of control untreated mice. Notably, the apparent mucosal healing effects of Sb576 is accompanied with reduced colon carcinogenesis, indicating that increase of luminal spermidine does not promote tumor development.

We propose that our engineered spermidine-secreting *S boulardii* represents a novel therapeutic targeting the greatest unmet challenge in current IBD management – mucosal repair and regeneration ^37^. Work is in progress to engineer *S boulardii* to secrete biologics that target key inflammatory molecules such as TNFα and the pro-regenerative spermidine as combinatorial therapeutics for IBD that might have the potential to break the current treatment ceiling ^69^.

## Materials and Methods

### Plasmid construction

All plasmids used in this study are listed in Supplementary table 2. For integration into the *Ty2* locus, DNA between the 5’ Ty2 and 3’ Ty2 homology sequences of pCfB2793 ^49^ was replaced by the coding regions of *SPE1*, *SPE2* or *TPO5* genes from *S. cerevisiae.* All genes were controlled by *TDH3* promoter and *CYC1* terminator and tagged with a FLAG epitope at the C-terminus. To reduce the complexity of the constructs, a fusion of the *SPE1* and *SPE2* reading frames was expressed from the same promoter with a 2A peptide cleavage site placed between the two reading frames ^70^. The *URA3* auxotrophic marker was replaced by *kanMX* to allow colony forming units (CFUs) to be determined by selection on G418 medium. The integration cassette was released from with *Not*I prior to transformation into *S. boulardii*.

### Yeast transformation

*S. boulardii* strains were transformed using the lithium acetate method ^71^. The wild type *S. boulardii* strain MYA-796 (ATCC) was inoculated and grown overnight at 30°C and 220 rpm. The overnight culture was diluted to an OD_600_ of 0.5 and incubated at 30°C and 220 rpm until it reached and OD_600_ of 1.5. Cells were then washed and 6.0 OD_600_ were resuspended in transformation mix (240 µL 50% PEG 3350, 36 µL 1M Lithium Acetate, 50 µL single-stranded carrier DNA (2 mg/mL in TE), 36 µL 1M DTT, and 16 µL water + DNA (1 µg DNA for *S. boulardii* and 100 ng DNA for *S. cerevisiae*). Cells were incubated at 42°C for 40 min, pelleted and washed with 1 mL sterile water, re-suspended in 200 µL water and immediately plated onto appropriate selective medium. For G418 selection, cells were recovered overnight in rich medium at 30°C and 220 rpm prior to plating.

### CRISPR-Cas9 mediated gene disruption

The plasmid pGZ110 expressing Cas9 and a targeting sgRNA was modified to replace the *URA3* auxotrophic selection marker with the *amdSYM* recyclable selectable marker ^72^. *amdSYM* confers resistance to acetamide and can be counter-selected using fluoroacetamide. The guides to target *OAZ1* were designed using the Synthego Knockout Guide Design tool (https://design.synthego.com/<u>#/</u>). The vector also contained a homology repair region that introduced several stop codons and a frameshift to prevent readthrough into the OAZ1 coding region, as well as a PAM site deletion to prevent continuous Cas9 nucleolytic activity. After the deletion was confirmed by Sanger sequencing, the plasmid was removed by counter-selection on fluoroacetamide medium.

### Animal procedures

All animal procedures have been approved by University of Ottawa Animal Care Committee. Mice were purchased from the Laboratories Charles River, Quebec, Canada (http://www.criver.com). All animals were kept at University of Ottawa’s animal care and veterinarian services facility, in a specialized pathogen-free environment (25 ± 2°C and 55% humidity, 12/12-hour light and dark cycle).

Yeast preparation, oral gavage, and determination of yeast cell viability in the gut: Yeast were cultured in YPD medium at 30 °C until saturation, harvested and washed twice in HEPES-saline resuspension buffer (100 mM HEPES, pH 7.0, 135mM NaCl). Washed yeast pellets were stored in single-use aliquots in HEPES-saline buffer containing 10% glycerol at -80 °C. After thawing, 200 μL of yeast suspension in buffer containing 10% glycerol was administered by oral gavage. To determine colony forming unit (CFU), fresh fecal samples were mixed with 20 × water (volume/weight) by vigorous vortexing. The samples were further serially diluted (10 ×) in water before plating on YPD yeast plates with ampicillin (100 μg /mL) or G418 (100 μg/mL), as indicated in legends. Plates were incubated at 30°C for 2 days followed by manual colony counting.

AOM-DSS chronic colitis and colon carcinogenesis: Mice were treated with a single dose of streptomycin (S9137; Sigma-Aldrich) via oral gavage (20mg in 200μL of PBS) on day -6, one day before the start of five (day -5 to day -1) daily oral gavages of yeast (1×10^8^ cells in 200 μL per mouse) or vehicle (200 μL of HEPES-saline+10% glycerol). On day 0, 16 hours following the fifth vehicle/yeast oral gavage, feces were collected for CFU and polyamine determinations. Mice then received a single injection of AOM (12.5mg/kg; i.p.; A5486, Sigma-Aldrich) on day 0 followed by three cycles of DSS treatment (2.5% in drinking water, 5 days for the first two cycles and 4 days for the third cycle) starting on day 5 with an interval of 21 days between cycles. Vehicle or yeast oral gavage continued three times a week after the first five doses. Daily disease activity index (DAI, consisting of weight loss, stool consistency, and the presence of blood in feces) scoring was performed according to Dong et, al. ^73^ with slight modifications.

Weight loss was measured against the body weight on day 5 (prior to DSS drinking water): none (0), 1%–5% (1), 6%–10% (2), 11%– 15% (3), >15% (4). Stool consistency was scored as normal (0), slightly loose (1), loose stool (2), pasty diarrhea (3), or watery diarrhea (4). Blood in stool was scored as negative, as assessed by fecal occult blood test kit (SK-61200, ThermoFisher) (0), occult blood test positive (2), and frank blood in feces or gross bleeding (4). Daily DAI for each mouse was the sum of the three scores. Weight changes and DAI scores were recorded for ten days following the start of each of the three DSS cycles. Mice were sacrificed 26 days following the end of the third DSS cycle for postmortem analyses.

### Tumor scoring, histopathology and immunohistochemistry

The entire colon was cut longitudinally, rinsed in PBS before being placed under a dissecting microscope for tumor measuring using a digital caliper. Immediately after, the colon was rolled into a Swiss roll with the distal end at the center, fixed in neutral buffered formalin and embedded in paraffin. A 5-micron section through the center of the Swiss roll was stained with Hematoxylin and eosin (H&E), dehydrated through an increasing concentration series of ethanol and xylene, and photographed using a slide scanner (ZEISS Axio Scan.Z1, Jena, Germany).

Morphologic features were assessed on coded images, according to criteria as previously described for humans ^55–57^ and mice ^40^. Briefly, low-grade adenomas were characterized by columnar epithelial cells depicting a high nuclear-to-cytoplasmic ratio, nuclear hyperchromasia and lack of surface maturation. With high-grade adenomas, over and above the features of low-grade adenomas, there was glandular complexity including irregularity of the architectural configuration. With progression to adenocarcinomas, there was fusion of glands to form a cribriform pattern, necrosis and a desmoplastic reaction.

Adjacent sections of the Swiss rolls were immunostained with anti-CD8 (Cell Signaling Technology, #98941, 1:100) or anti-Granzyme B (Cell Signaling Technology, #44153, 1:100) and counterstained with Hematoxylin, done by Louise Pelletier Histology Core Facility of University of Ottawa. The slides were imaged using a Zeiss AxioScanner. Coded images were examined to identify reactive lymphoid follicles (RLF) and neoplastic lesions (as above). The proportion of CD8^+^ (or Granzyme B+) cells within RLFs or neoplastic lesions were quantified using QuPath and expressed as % of the total nuclei there. The % CD8^+^ (or GranzB+) of each mouse was weighted average of all RLFs (or neoplastic lesions) of the mouse.

### Polyamine determination

Intracellular yeast polyamines were extracted according to Qin, et al. ^48^. One mL saturated yeast culture was centrifuged (2000 rpm, 5 min) and the pellets resuspended in 495 μL of boiling water plus 5 μL of 100 μg/mL deuterated spermidine, Spermidine-(butyl-d_8_) trihydrochloride (709891; Sigma-Aldrich). The tube was then vortexed vigorously and kept at 100°C on a heat block for 30 min. The post centrifugation (twice at 15,000 rpm at 4°C,15 min) supernatant was then stored at –20 °C or directly subjected to the quantification step.

Yeast extracellular (medium) polyamines and free fecal polyamines were extracted using BondElut Plexa cartridges (12113001; Agilent Technologies). For medium samples, tubes containing 1 mL saturated yeast culture were centrifuged (2000 rpm, 5 min) and the supernatant used for the next steps. BondElut Plexa cartridges were placed on a vacuum apparatus (Agilent Technologies). After washing the cartridges with 500μl methanol and 500μL ddH_2_O, 550.8μL of media sample including 500μl of saturated yeast media + 5μL of 10μg/mL Spermidine-(butyl-d_8_) trihydrochloride and 45.8 μL of 18% NH_4_OH (AX1303; Sigma-Aldrich) were applied to the cartridge. After washing the cartridges with 500μL 5% methanol/ddH_2_O, polyamines were eluted with 500μL of 2% formic acid/methanol. The elutes were then stored at –20 °C or directly subjected to the quantification step.

To extract free fecal polyamines, 50-100 mg of feces was mixed with 10 volumes of PBS, vortexed vigorously for 10 sec for 3 times, placed on ice for 5 min, and then centrifuged at 14000 rpm for 15 min. 500 μL of the supernatant was mixed with 50μL of Spermidine-(butyl-d_8_) trihydrochloride and 50μL of 18% NH_4_OH. The mixture was centrifuged at 14000 rpm for 10 min and passed through BondElut Plexa cartridges as described above. The eluates were then stored at –20 °C or directly subjected to the quantification step.

All extracted samples were analyzed by using a validated liquid chromatography mass spectrometry/mass spectrometry (LC-MS/MS) method. The chromatographic separation was achieved on the Thermo Accela LC system with a Supelcosil AZB+Plus 3μm column (100×2.1mm). Gradient elution (0.1% heptafluorobutyric acid as the ion-pairing agent with acetonitrile, 300μl/ml at 40°C) was used to separate all polyamines and internal standard (Spermidine-(butyl-d_8_) trihydrochloride). Mass spectral analyses were accomplished on a quadrupole tandem Mass Spectrometry (TSQ Quantum Access MAX, Thermo). Nitrogen was used as curtain, nebulizer and collision gases. User controlled voltages, gas pressures, and source temperature was all optimized for the detection of polyamines and IS. MS was quantified using electrospray multiple reaction monitoring (MRM) in positive mode and the MRM transitions were m/z 89.1to 72.1 for putrescine, m/z 146.2 to 112.2 for spermidine m/z 203.2 to 129.1 for spermine, and m/z 154.1 to 120.3 for IS. 10 μL was used for injection and the autosampler tray temperature was 22 °C. The effective linear range was 0.2 -5μg/mL for putrescine, 01-2.5µg/mL for spermidine, and 0,05-1 μg/mL for spermine. Inter-batch precision (CV%) varied between 6 and 18% and intra-batch accuracy varied between 87 and 118%. Validation results displayed that all polyamines and IS were stable for at least 10 hours at 22 °C. Intracellular polyamine concentrations were determined assuming *S. boulardii* cell volume as 33μm^3^. Published data of intracellular concentration for *S cerevisiae* is 1.34mM ^74^.

### Statistical Analysis

Statistical analyzing and graphing were performed using GraphPad Prism software version 9 (Boston, MA, USA). Quantitative variables were reported as means ± standard error of the mean (SEM). The difference of means was analyzed by One-way ANOVA and Tukey’s multiple comparisons test. *: p<0.05; **: p<0.01; ***: p<0.001; ****: p<0.0001.

