## supplemental tables for "Engineered spermidine-secreting *Saccharomyces boulardii* ameliorate colitis and colon cancer in mice"

Supplementary Table 1. Yeast strains used in this study.

| Name | Number | Relevant Genotype | Source |
| --- | --- | --- | --- |
| SbWT | MTy5073 | MYA-796 | ATCC |
| Sb570 | MTy5331 | MYA-796 *oaz1Δ TY2::pTDH3-TPO5-FLAG pTDH3-SPE1-FLAG-KanMX::Ty2* | This study |
| Sb576 | MTy5532 | MYA-796 *oaz1Δ TY2::pTDH3-TPO5-FLAG pTDH3-SPE1-2A-SPE2-FLAG-KanMX::Ty2* | This study |

Supplementary Table 2. Plasmids used in this study.

| Name | Number | Construction | Source |
| --- | --- | --- | --- |
| OAZ1 CRISPR knockout | pMT4951 | pGZ110-Cas9-sgRNA oaz1Δ-amdSYM; OAZ1-sgRNA (5’-ATAGAGGACCCATCAAATCT-3’) cloned into pGZ110 base vector (gift from B. Futcher) | This study |
| TPO5/SPE1 integration cassette | pMT4952 | pcfb2974-NotI-Ty2-pTDH3-TPO5-FLAG-pTDH3-SPE1-FLAG-KanMX-Ty2-NotI | This study |
| TPO5/SPE1/SPE2 integration cassette | pMT4953 | pcfb2974-NotI-Ty2-pTDH3-TPO5-FLAG-pTDH3-SPE1-2A-SPE2-FLAG-KanMX-Ty2-NotI | This study |
| pKS-ST 2µm expression vector | pMT4954 | pKS-pADH2-FLAG-TCyc1-KanMX-2µm | Dual System Biotech |
